# Replicator dynamics generalized for evolutionary matrix games under time constraints

**DOI:** 10.1101/2024.08.22.609164

**Authors:** Tamás Varga

## Abstract

One of the central results of evolutionary matrix games is that a state corresponding to an evolutionarily stable strategy (ESS) is an asymptotically stable equilibrium point of the standard replicator dynamics. This relationship is crucial because it simplifies the analysis of dynamic phenomena through static inequalities. Recently, as an extension of classical evolutionary matrix games, matrix games under time constraints have been introduced (Garay et al. 2017; Křivan and Cressman 2017). In this model, after an interaction, players do not only receive a payoff but must also wait a certain time depending on their strategy before engaging in another interaction. This waiting period can significantly impact evolutionary outcomes. We found that while the aforementioned classical relationship holds for two-dimensional strategies in this model, it generally does not apply for three-dimensional strategies (Varga and Garay 2024). To resolve this problem, we propose a generalization of the replicator dynamics that considers only individuals in active state, i.e., those not waiting, can interact and gain resources. We prove that using this generalized dynamics, the classical relationship holds true for matrix games under time constraints in any dimension: a state corresponding to an ESS is asymptotically stable. We believe this generalized replicator dynamics is more naturally aligned with the game theoretical model under time constraints than the classical form. It is important to note that this generalization reduces to the original replicator dynamics for classical matrix games.

**Mathematics Subject Classification (2020):** 37N25, 91A05, 91A22, 91A40, 92D15, 92D25

## 1 Introduction

Recently, some studies discussed the incorporation of time constraints into evolutionary matrix games (Garay et al. 2017; Křivan and Cressman 2017). This concept is rooted in Holling’s functional response (Holling 1959a; Holling 1959b), which considers the temporary inactivity of a predator following a successful hunt. Motivated by this, matrix games under time constraints model pairwise interactions with dual consequences: dependent on the strategy of each player, they receive some payoff and have to wait for a certain amount of time before becoming ready for another interaction. For instance, the payoff could represent food or injury, while the corresponding waiting time reflects the time needed for handling/digesting food or recovering from an injury. Both the payoff and the waiting time depend on the strategy of the focal individual and its opponent. It is important to emphasize that these two factors collectively determine a player’s success because if a player has to wait a substantial amount of time after interactions, the average income per unit of time may be low even if the payoffs following the interactions are high.

Evolutionary game theory fundamentally adopts a dynamical perspective, although the initial mathematical formulation of evolutionary stable strategy emerged in a static way (Maynard Smith and Price 1973; Maynard Smith 1974). Propelled by the dynamic aspect, Taylor and Jonker (1978) introduced the replicator dynamics (also known as the replicator equation) to model the evolution of a polymorphic population, establishing a connection between the evolutionary stability of a strategy and the asymptotic stability of the corresponding point in the replicator dynamics. This linkage, implying that the former guarantees the latter, is a central tenet of evolutionary game theory (Hofbauer et al. 1979; Zeeman 1980).

Our recent analysis explored analogous relationships in the presence of time constraints within the model. It was validated that the state corresponding to an evolutionarily stable strategy remains an asymptotically stable equilibrium point of the replicator dynamics, assuming the strategies are two-dimensional, meaning each strategy is a mixture of the same two pure strategies (Garay et al. 2018). In higher dimensions, however, we were only able to establish this implication in specific cases, such as when the evolutionarily stable strategy is a pure strategy (Varga et al. 2020). Our conjecture was that this generalization is not universally true, and indeed, counterexamples were successfully constructed (Varga and Garay 2021, Varga and Garay 2024).

In this article, we present a generalization of replicator dynamics that accounts for time constraints. Broadly speaking, the fundamental idea behind this generalization is that an inactive (waiting) individual cannot interact and therefore does not participate in resource allocation through interactions. Since, in the model, the sole difference in fitness among different strategies arises from their performance in interactions, the success of a given type depends on its proportion among the active individuals rather than its proportion in the entire population. This suggests that fitness should be interpreted as an amount proportional to the rate of change in frequency among active individuals rather than the frequency with respect to the whole population. Following this idea replicator dynamics can be generalized in such a way that it is in harmony with the game theoretical model in the sense that the state corresponding to an evolutionarily stable strategy is an asymptotically stable equilibrium point of the generalized dynamics with respect to the pure strategies. It is worth noting that this is a genuine generalization, as it simplifies to the usual replicator dynamics in the case of classical evolutionary matrix games (the particular case, when time constraints are independent of the strategies). Moreover, from a biological viewpoint, it continues to be replicator dynamics since an individual of type *i* can produce and be descended from only individuals of type *i*.

## 2 Preliminaries

Here, we review the mathematical formalism and previous results necessary for the discussion of the new findings. The detailed setup, the mathematical derivation of the formulas and possible interpretation of the model adhere to those presented in Garay et al. (2017), Garay et al. (2018) and Varga et al. (2020). We envision a vast population, with the number of individuals approaching infinity, engaging in random pairwise interactions. These interactions yield two outcomes for each participant, depending on their strategies. One outcome is the conventional payoff, akin to classical matrix games, such as obtaining food or suffering injury during the interaction. In addition to the payoff, there is another consequence: the **waiting time** that a player must endure after the interaction. We assume that the waiting time follows an exponential distribution, and its expected value depends not only on the strategy used by the focal player but also on the strategy used by the opponent. During this period, the individual relaxes and refrains from seeking a new opponent or engaging in further interactions. Once the waiting time elapses, the individual resumes the search for a new opponent to initiate another interaction. We remark that if the waiting time is independent of the players’ strategies, meaning it has the same duration after each interaction, the scenario reverts to classical evolutionary matrix games, solely considering the payoffs resulting from interactions (Maynard Smith 1974, Chapter 2 in Maynard Smith 1982, or Chapter 6 in Hofbauer and Sigmund 1998).

If an individual is searching for an opponent, we say that the individual is **active**. Active in the sense that, when it encounters another individual also in search of an opponent, they can engage in interaction with each other. We also assume that the time required for a search follows an exponential distribution, but its expected value is independent of the players’ strategies. We can therefore consider its value as the unit of time. If the searching individual encounters a waiting individual, no interaction occurs. This type of encounter has no consequences for the waiting individual in the mathematical model; they simply continue waiting as if nothing happened. The searching individual, on the other hand, initiates a new search. Therefore, the more individuals are inactive, the longer it takes to find another active individual. We refer to an individual that is in a waiting period following an interaction as **inactive**, since, as mentioned above, a waiting individual is not capable of interaction. The time necessary for the interaction itself can be assumed to be 0 in the model or can be considered as part of the waiting time. We assume that the population itself is in a stationary state at every moment. This means that if the frequencies of the admissible phenotypes are fixed, the proportion of active and inactive individuals respectively remains constant, even within subpopulations of different phenotypes: the number of individuals becoming inactive equals the number becoming active. For more technical details about the model, refer to Garay et al. (2017).

An individual has the option to adopt either a pure strategy or a combination of admissible pure strategies. When an individual employing pure strategy *i* engages in an interaction with another individual using pure strategy *j*, the payoff for the individual choosing pure strategy *i* is denoted by *a*_*ij*_, and the waiting time for this individual after the interaction is *τ*_*ij*_. The value of *a*_*ij*_ can be negative, zero, or positive, while *τ*_*ij*_ is assumed to be non-negative. If there are *N* admissible pure strategies, the *N×N* matrix composed of elements *a*_*ij*_ is termed the **payoff matrix**, and the *N×N* matrix composed of elements *τ*_*ij*_ is referred to as the **time-constraint** matrix. Formally, the strategies precisely constitute the elements of the *N* -dimensional standard simplex:

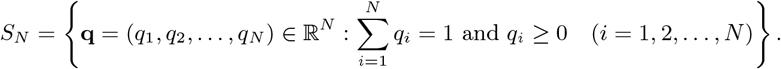

The **pure strategies** are strategies in which only one coordinate is 1, and the others are 0. We denote by **e**_*i*_ the pure strategy, with the *i*-th coordinate being 1 and the other coordinates being 0. Clearly, there are *N* pure strategies in *S*_*N*_. Strategies distinct from the pure strategies are called **mixed strategies**. Any arbitrary strategy **q** can be interpreted as a probability distribution, giving the probability that an individual with strategy **q** uses the *i*-th pure strategy in an interaction, namely, this probability is *q*_*i*_. We remark that the terms “phenotype” or simply “type” are often used interchangeably with the term “strategy”. Additionally, we use the phrase “**q** individual/strategist/phenotype” to refer to an individual following strategy **q**.

In the following, we will review the polymorphic and monomorphic approaches. In the polymorphic approach, we consider a population in which a fixed, finite number of phenotypes can occur, and the focus is not on the strategy of a specific individual but on the average strategy of the population. In the monomorphic approach, we examine a population where every individual belongs to a specific phenotype, the resident phenotype, and we study what happens evolutionarily if a small proportion of another so-called mutant phenotype appears. This other phenotype, different from the resident strategy, can be any strategy, but at most one mutant strategy is allowed at a time. The monomorphic approach is essential for defining the concept of evolutionary stability, while the polymorphic approach is necessary for describing the temporal changes in a population.

First, we will discuss the polymorphic approach, allowing for the presence of an arbitrary number (though finite and pre-fixed) of phenotypes in the population. Consider a population where the possible phenotypes of individuals are **f**_1_, **f**_2_, … **f**_*n*_. No other phenotypes are possible besides these. The frequencies of individuals associated with each phenotype are denoted by *x*_1_, *x*_2_, …, *x*_*n*_, respectively (*x*_*i*_ ≥ 0, *i* = 1, 2, …, *n* and ∑*i x*_*i*_ = 1). Then there is a unique solution (*ϱ*_1_, *ϱ*_2_, …, *ϱ*_*n*_) ∈ [0, 1]^*n*^ of the equation system

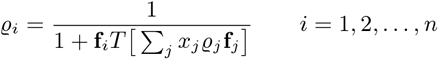

(Lemma 2. in Garay et al. 2017)^1^. Where it is important to highlight the dependence of the solution on the frequency distribution **x** = (*x*_1_, *x*_2_, …, *x*_*n*_), we use the notation *ϱ*_*i*_(**x**). *ϱ*_*i*_ can be considered as the proportion of active individuals within the subpopulation of individuals following strategy **f**_*i*_. In the equations, the numerator 1 represents the expected time required to find another individual (active or inactive) from the end of the waiting time after the last interaction. As previously mentioned, it is set to 1 as this duration is considered the unit of time. When an individual searching for an opponent encounters an inactive individual, no interaction takes place. This scenario is treated as an interaction with zero intake and zero waiting time, after which the search recommences. Therefore, the expression **f**_*i*_ *T* [∑_*j*_*x*_*j*_ *ϱ*_*j*_ **f**_*j*_ ] describes the average waiting time following an interaction, while 1 + **f**_*i*_*T*[∑_*j*_ *x*_*j*_*ϱ*_*j*_ **f**_*j*_ ]represents the average time between two successive encounters.

Similarly, since an active individual encounters another active individual with probability ∑_*i*_ *x*_*i*_*ϱ*_*i*_ and the intake is 0 if the opponent is inactive, **f**_*i*_*A* [∑_*j*_ *x*_*j*_*ϱ*_*j*_ **f**_*j*_ ] is the expected intake in an interaction.

Then the **fitness** of an **f**_*i*_ individual is defined as the following intake rate:

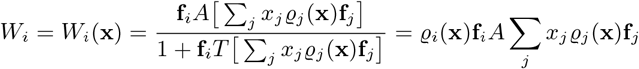

Later we use the next notations:

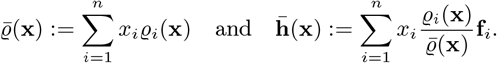

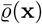 gives the proportion of the active individuals with respect to the whole population while 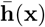 is the average strategy of the active individuals.

To define evolutionary stability, we require the monomorphic setup. This approach, initially employed by Maynard-Smith and Price in their pioneering article (Maynard Smith and Price 1973), involves a vast population where nearly all individuals adopt the same strategy, known as the resident strategy. The remaining portion of the population adheres to a different strategy, termed the mutant strategy. At any given moment, there is at most one mutant strategy, but it can be any strategy from the strategy space distinct from the resident one. If the resident strategy is such that its fitness is greater than that of any mutant phenotype providing the proportion of mutants are small enough then the resident strategy is called evolutionary stable.

To precisely define evolutionary stability for matrix games under time constraints, we introduce analogues of the polymorphic setup. Let **p** and **q** be strategies and consider a population consisting of **p** and **q** individuals with frequencies (1−*ε*) and *ε*, respectively. Then *ρ*_**p**_ = *ρ*_**p**_(**p, q**, *ε*) denotes the proportion of active individuals in the subpopulation of **p** individuals. Similarly, *ρ*_**q**_ denotes the proportion of active individuals in the subpopulation of **q** individuals. The values of *ρ*_**p**_ and *ρ*_**q**_ are calculated as the unique solution of the subsequent equation system:

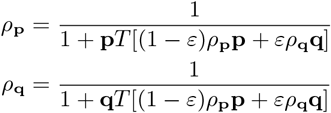

(Lemma 2. in Garay et al. 2017). Then the fitness of a **p** individual is

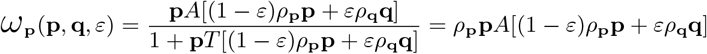

and that of **q** individual is

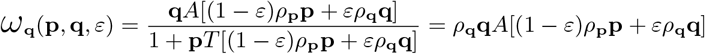

When *ε* = 0 we simply write *ρ*(**p**), *ρ*_**q**_(**p**), *ω*(**p**) and *ω*_**q**_(**p**), respectively, instead of *ρ*_**p**_(**p, q**, 0), *ρ*_**q**_(**p, q**, 0), *ω*_**p**_(**p, q**, 0) and *ω*_**q**_(**p, q**, 0), respectively. Using the notations above, the evolutionary stability can be formulated in our model as follows.

### Definition 2.1

*The strategy* **p** *is said to be a* ***uniformly evolutionarily stable strategy*** *(UESS for short) if there is an ε*_0_ *>* 0 *such that, for any* **q***≠* **p** *and* 0 *< ε < ε*_0_, *we have that*

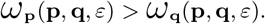

We also need the following characterization based on neighbourhood invader property (cf. Apaloo 1997; Apaloo 2006, Theorem 6.4.1 in Hofbauer and Sigmund 1998).

### Lemma 2.2

**(Varga et al. 2020, Theorem 3.2)** *A strategy* **p** *is a UESS if and only if there is δ such that*

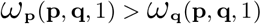

*whenever* 0 *<* ||**p** − **q**|| *< δ*.

Using the definition of *ω*_**p**_ and *ω*_**q**_, the above inequality can be write as *ρ*_**p**_(**q**)**p***Aρ*(**q**)**q** *> ρ*(**q**)**q***Aρ*(**q**)**q** which is obviously equivalent to the inequality *ρ*_**p**_(**q**)**p***A***q** *> ρ*(**q**)**q***A***q**.

The adverb “uniformly” refers to the independence of *ε*_0_ from **q**. In Maynard-Smith and Price’s classical model, this independence is not assumed (Maynard Smith and Price 1973; Taylor and Jonker 1978), but it can be proven (Theorem 6.4.1 in Hofbauer and Sigmund 1998). For our model, however, it is an open question, whether the uniformity can be derived if *ε*_0_ can vary with **q**, that is, when **p** is only assumed to be an ESS rather than a UESS. It would not be surprising if there is no equivalence because the fitness functions *ω*_**p**_ and *ω*_**q**_ are generally non-linear in both **p** and **q**. This is in contrast to the classical model which can be considered as a special case when every element of *T* is the same.

The definition of ESS aims to express the expectation that an evolutionarily stable strategy can resist any type of mutants, in the sense that if the proportion of mutants is small enough then the mutant strategy will be eliminated by natural selection from the population. This dynamical aspect of ESS is not directly present in the above definition but if we consider the dynamics

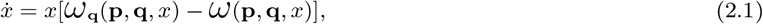

where *ω*(**p, q**, *x*) = (1−*x*)*ω*_**p**_ + *xω*_**q**_ and *x* is the frequency of the **q** individuals, then it can be easily seen that **p** is an ESS if and only if the state *x* = 0 is an asymptotically stable rest point of the previous dynamics for any **q**≠ **p** (c.f. the reasoning around dynamics (7.14) on p. 72 in Hofbauer and Sigmund 1998). To ensure the uniformity, it should also be demanded the existence an *ε*_0_ *>* 0 such that if *x*(0) *< ε*_0_ then *x*(*t*) *→* 0 under dynamics (2.1) for any **q** ≠ **p**.

For every **q** ≠ **p**, the dynamics described above is just a replicator dynamics which says that the change in the frequency of a particular type in a population is proportional to both the frequency and the relative fitness of that type. This dynamics was first suggested by Taylor and Jonker (1978). In the polymorphic case it takes the following shape

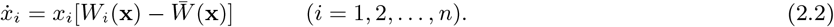

This is the replicator dynamics (or replicator equation) with respect to the apriori fixed phenotypes **f**_1_, **f**_2_, …, **f**_*n*_ where *x*_*i*_ is the frequency of **f**_*i*_ individuals. In this article we focus on **the standard replicator dynamics**, that is, when *n* = *N* and **f**_*i*_ = **e**_*i*_.

One of the fundamental theorems of the theory of classical evolutionary matrix games (that is, when every element of *T* is the same) claims that if **p** is a UESS then state **x** = **p** = (*p*_1_, *p*_2_, …, *p*_*N*_) is an asymptotically stable rest point of the standard replicator dynamics (Hofbauer et al. 1979; Zeeman 1980; Theorem 7.2.4 in Hofbauer and Sigmund 1998).

To extend this theorem for arbitrary matrix games under time constraints we should first consider the corresponding state of a strategy.

### Lemma 2.3

**(Garay et al. 2018, Proposition 3.1 and Corollary 6.3)** *Consider the polymorphic population of* **e**_1_, **e**_2_,…, **e**_*N*_ *individuals in state* **x** *and let*

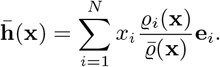

*Then* 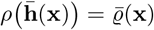 *and* 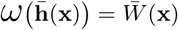.

*Furthermore, for any strategy* **p** ∈ *S*_*N*_, *there is a unique state* **x** *such that* 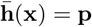, *namely*, 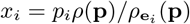.

The lemma says that if one consider the monomorhic population of 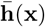 individuals then the fitness of an individual in this population agrees with the fitness of an “average” active individual in the polymorphic population.

The state **x** with 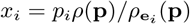 in the above lemma is denoted by **x**(**p**).

### Definition 2.4

*Strategy* **p** *is a* **Nash equilibrium** *for the matrix game under time constraints* (*NE for brevity*), *if for all* **q***≠* **p** *we have*

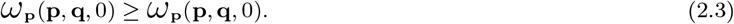

*If the above inequality is strict for every* **q***≠* **p** *then* **p** *is called a* ***strict Nash equilibrium***.

Note that the above inequality is equivalent to the inequality *ϱ*(**p**)**p***A***p** ≥*ϱ*_**p**_(**q**)**q***A***p** (just simplify by *ϱ*(**p**)).

If strategy **p** is a Nash equilibrium, then **x**(**p**) is a rest point of the standard replicator dynamics showing that **x**(**p**) takes over the role of state **p** under time constraints (Lemma 3.2 in Garay et al. 2018). Since a UESS is a Nash equilibrium^2^, it follows that if a strategy **p** is a UESS then the corresponding state **x**(**p**) is a rest point of the replicator dynamics. But what about the stability of this rest point? As mentioned above, the rest point is asymptotically stable in the case of classical evolutionary matrix games (when elements of T are identical). However, except for the case of *N* = 2 (Theorem 4.2 in Garay et al. 2018), this is generally not true. Even for *N* = 3, there are examples where the corresponding rest point **x**(**p**) of the UESS **p** is an unstable rest point of the replicator dynamics (Varga and Garay 2021, Varga and Garay 2024). This is particularly unfavorable from a biological perspective because the condition formulated in Definition 2.1 regarding evolutionary stability is not sufficient to imply stability in a dynamic model. In this article, we propose a generalization of replicator dynamics that takes into account time constraints, and the equilibrium point corresponding to a UESS is asymptotically stable. Moreover, it simplifies to the replicator dynamics in the case of classical evolutionary matrix games.

## 3 Generalization of the replicator dynamics

To establish a relationship between polymorphic and monomorphic approach the mean strategy of active individuals of the polymorphic population should be considered according to Lemma 2.3 rather than the mean strategy of all the individuals of the polymorphic population. This suggests that the fate of the polymorphic population primarily depends on the composition of the subpopulation of active individuals. Investigate this phenomenon from a mathematical perspective.

We first recall the classical evolutionary matrix games, that is, when all elements of *T* are the same and so 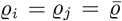for every *i, j* ∈ {1, 2, …, *N* }. Assume that **p** is an ESS for this game^3^. Then there is a neighbourhood ℬof **p** = (*p*_1_, …, *p*_*N*_) such that

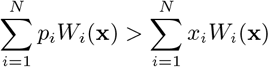

whenever **x** ∈ *ℬ* \ {**p**} (see Chapter 6.4, 7.1 and 7.2 Hofbauer and Sigmund 1998). In other words, being ESS implies that **p** is a polymorphic stable state^4^ or PS state for short (now **p** =∑*p*_*i*_**e**_*i*_). This intuitively means that if state **x** is close enough to state **p** then a subpopulation in state **p** has a higher average fitness in a population in state **x** than the average fitness of the whole population. Therefore one can expect that the population evolves toward state **p** in the “next time unit”.

Indeed, if **p** is an ESS then **p** is a PS state and so an assymptically stable rest point of the replicator dynamics (Hofbauer et al. 1979; Zeeman 1980; Theorem 7.2.4 in Hofbauer and Sigmund 1998).

However, this connection generally holds no longer in present of time constraints. Though, by Lemma 3.2 in Garay et al. (2018), we know that **x**(**p**) is a rest point of the replicator dynamics, examples in Varga and Garay (2021) and Varga and Garay (2024) show that state **x**(**p**) can be unstable. Hence, the (U)ESS property can not imply the PS property of the corresponding state in general.

In the case of classical evolutionary matrix games, the “*ϱ*” and “*ρ*” functions do not depend on the composition of the population, the strategies and frequencies, and they are just constant positive multipliers which can be omitted. This implies that 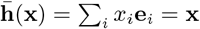 holds for every **x** ∈ *S*_*N*_. Whereas, under time constraints, it is not at all certain that 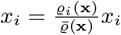 or 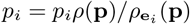.

Let 𝒴 (**x**) be the set of states **y** (distinct from **x**) such that the average fitness of a subpopulation in state **y** of a population in state **x** is greater than the average fitness of the population, that is,

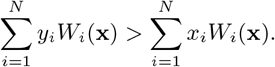

As mentioned previously, in the case of classical matrix games, if **x** close enough to **p**, then **p** ∈ *𝒴*(**x**) (Theorem 6.4.1 in Hofbauer and Sigmund 1998). What can we state under time constraints? Let

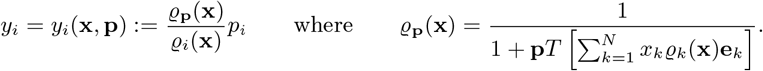

It is clear that *y*_*i*_ ≥ 0, furthermore,

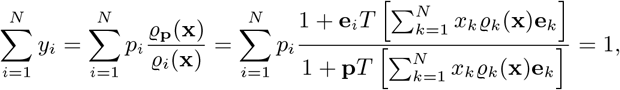

that is, (*y*_1_, …, *y*_*N*_) ∈ *SN*. Since 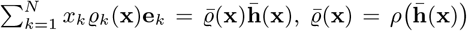 and 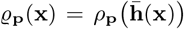 (see Lemma 2.3), it follows that

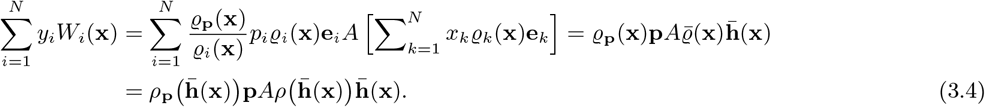

The rightmost side is continuous in **x** (Corollary 6.3 in Garay et al. 2018). Therefore, if **p** is a UESS then there is an *η >* 0 such that

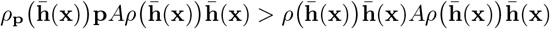

holds for any **x** ∈ *S*_*N*_ with 0 *<*|| **x** −**x**(**p**) ||*< η* (use the characterization of UESS in Lemma 2.2). Consequently, (3.4) goes on as

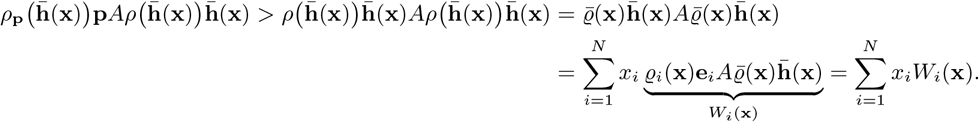

We summerize the previous reasoning in the following lemma.

### Lemma 3.1

*Assume that* **p** *is a UESS. Let*

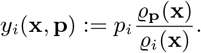

*Then there exists an η >* 0 *such that*

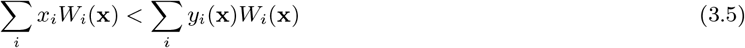

*whenever* 0 *<* ||**x** − **x**(**p**)|| *< η*.

In other words we have got that the relation *y*_1_(**x, p**), …, *y*_*N*_ (**x, p**) ∈ 𝒴(**x**) under time constraints is a generalization of the relation **p** ∈ *𝒴*(**x**) in the classical case.

The previous observations leads us to introduce the following dynamics:

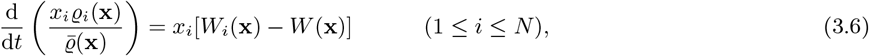

where 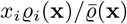 on the left-hand side is the frequency of the *i*-th pure phenotype in the subpopulation of active individuals. This frequency is just the *i*-th coordinate of the mean strategy of active individuals which represents the population by Lemma 2.3. The equation says that the rate of change in the frequency of the active **e**_*i*_ individuals among the active individuals is proportional to both the frequency and the relative fitness of the *i*-th phenotype. According to this model, the evolution of the *i*-th phenotype occurs directly through the active subpopulation of the *i*-th phenotype. The intuitive explanation for this may be that only active individuals are capable of interaction, and thus the distribution of resources depends solely on them. Therefore, the success of a phenotype is better reflected by the change in the ratio among active individuals rather than the change in the ratio for the entire population. However, the right side remains the same as in the classical case, that is, the product of relative fitness and the frequency with respect to the entire population, although, at the first moment, one might expect the frequency among active individuals here as well. Nevertheless, this is not the case, as the model does not distinguish between active and inactive individuals in terms of reproduction. Whether an individual is active or inactive only determines its ability to interact, while the income from interactions is used equally by all individuals for reproduction, regardless of whether they are active or inactive. Note that, for classical evolutionary matrix games, dynamics (3.6) coincides with the standard replicator dynamics.

Observe that, by Lemma 2.3 dynamics (3.6) can be rewritten in the following form:

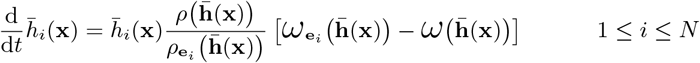

where 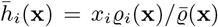 is the *i*-th coordinate of 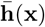, so introducing the variables 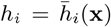 we can see that dynamics (3.6) is equivalent to the following dynamics:

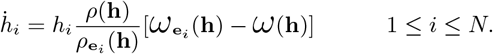

or, if the notation **x**(**p**) (introduced in the paragraph above Definition 2.4) is used,

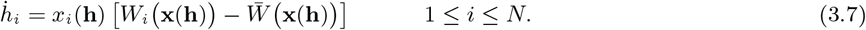

Here, *h*_*i*_ can be interpreted as the proportion of **e**_*i*_ individuals among the active individuals in the polymorphic population. *x*_*i*_(**h**) gives the proportion of **e**_*i*_ individuals as a function of **h**. Also, *W*_*i*_(**x**(**h**)) is the fitness of an **e**_*i*_ individual while 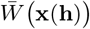 is the average fitness as functions of **h**.

Since *ρ*(**p**) is defined as the solution lying between 0 and 1 of the equation

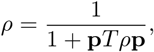

one can easily check that

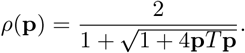

Hence, the right-hand side of dynamics (3.7) is continuously differentiable ensuring the existence and uniqueness of the solution of initial value problems. Also, since 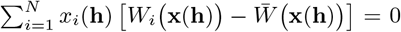 it follows, that the simplex *S*_*N*_ invariant under dynamics (3.7).

The relationship between a NE **p** and the corresponding state (which is **p** itself) of dynamics (3.7) is similar to that in classical evolutionary matrix games (cf. Theorem 7.2.1 in Hofbauer and Sigmund 1998 or Lemma 4.6 in Varga et al. 2020). int*S*_*N*_ below denotes the interior of *S*_*N*_, that is, the set of strategies with no zero coordinate.

### Lemma 3.2

a. *If* **p** *is a NE of the matrix game under time constraints then* **p** *is a rest point of dynamics (3.7)*.
b. *If* **p** ∈ int*S*_*N*_ *is a rest point of dynamics (3.7) then* **p** *is a NE of the matrix game under time constraints*.
c. *If* **p** *is a stable rest point of dynamics (3.7) then* **p** *is a NE of the matrix game under time constraints*.
d. *Assume that the singleton* {**p**}⊂*S*_*N*_ *is the ω-limit of an orbit* **h**(*t*) *running in* int*S*_*N*_ *under dynamics (3.7). Then* **p** *is a NE of the matrix game under time constraints*.

*Proof*. The proof follows the way of the proof of Theorem 7.2.1 in Hofbauer and Sigmund (1998) or Lemma 4.6 in Varga et al. (2020). Note that 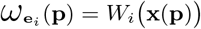 and 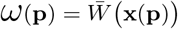.

a. Since **p** is a NE, it follows that 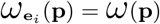 whenever *p*_*i*_*≠* 0 (Lemma 4.6 in Varga et al. 2020). Therefore,if *p*_*i*_*≠* 0 then *W*_*i*_ **x**(**p**) 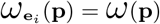. If *p*_*i*_ = 0 then *x*_*i*_(**p**) = 0. Hence 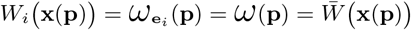 0 for every *i* ∈ *{*1, 2, …, *N}*, that is, **p** is a rest point of dynamics (3.7).
b. Since **p** is a rest point of dynamics (3.7), it follows that 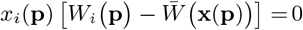 for every *i* ∈ *{*1, 2, …, *N}*. As **p** ∈ int*S*_*N*_, that is, *p*_*i*_*≠*0 and so *x*_*i*_(**p**)*≠*0 (Lemma 6.2 in Garay et al. 2018) for every *i* ∈ *{*1, 2, …, *N}*, this is possible if and only if 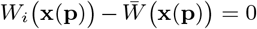. Hence, 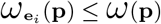, which means, by Lemma 2.3 in Varga et al. (2020) that **p** is a NE.
c. If *p*_*i*_*≠* 0 then, following point (b), we get that 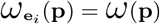. If *p*_*i*_ = 0 and 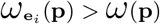 then, by continuity, there is a *κ >* 0 and a bounded neighbourhood 𝒩of such that 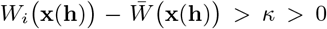 whenever **h** ∈ *𝒩* ∩ *S*_*N*_. Since **p** is also assumed to be stable, it follows the existence of another neigbourhood 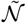 such that **h**(*t*) ∈ *N* ∩ *S*_*N*_ for every *t >* 0 if 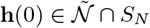. If 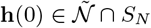 with *h*_*i*_(0) *>* 0 then

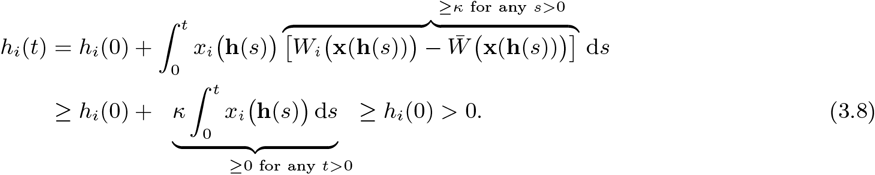 Using this lower estimate we get that

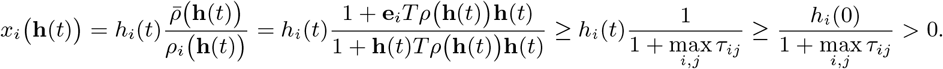

for every *t* ≥ 0. Hence

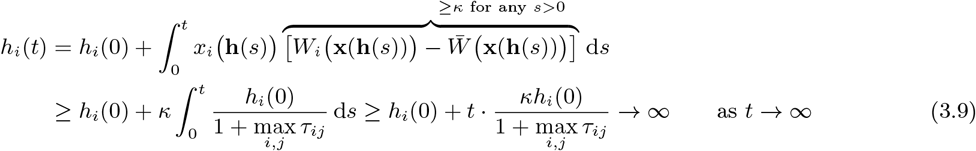

which contradicts the fact that **h**(*t*) ∈ *𝒩* ∩ *S*_*N*_ for every *t >* 0. So 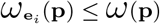 has to hold even if *p*_*i*_ = 0.
d. Assume that 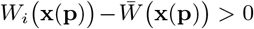 for some *i*. Then, by continuity, there is a *t*_0_ and *κ* such that 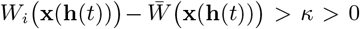 if *t* ≥ *t*_0_. We can set *t*_0_ = 0. As **h**(*t*) is assumed to be in int*S*_*N*_ we have that *h*_*i*_(0) *>* 0. Now, just repeat (3.8) and (3.9) to obtain a contradiction.

We have arrived at the analogue of Hofbauer et al. (1979), Theorem 1 in Zeeman (1980) or Theorem 7.2.4 in Hofbauer and Sigmund (1998).

### Theorem 3.3

*Assume that* **p** ∈ *S*_*N*_ *is an UEES then state* **p** *is a locally asymptotically stable rest point of dynamics (3.7)*.

*Proof*. From the previous lemma, it immediately follows that **p** is a rest point of dynamics (3.7). (To see that **p** is a NE, just let *ε* tend to zero in Definition 2.1.)

From here, the proof follows the proof of Theorem 7.2.4 in Hofbauer and Sigmund (1998). We verify that the function 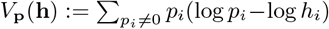 is a Lyapunov function with respect to dynamics (3.7). Since 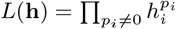 attains its strict maximum on *S*_*N*_ at *h*_*i*_ = *p*_*i*_ (*i* = 1, 2, …, *N*) (see the function *P* (**x**) in the proof of Theorem 7.2.4 in Hofbauer and Sigmund (1998)), itt follows that *V*_**p**_(**h**) is positive if **h** ≠ **p** and 0 at **h** = **p**, that is, positive definite.

On the other hand,

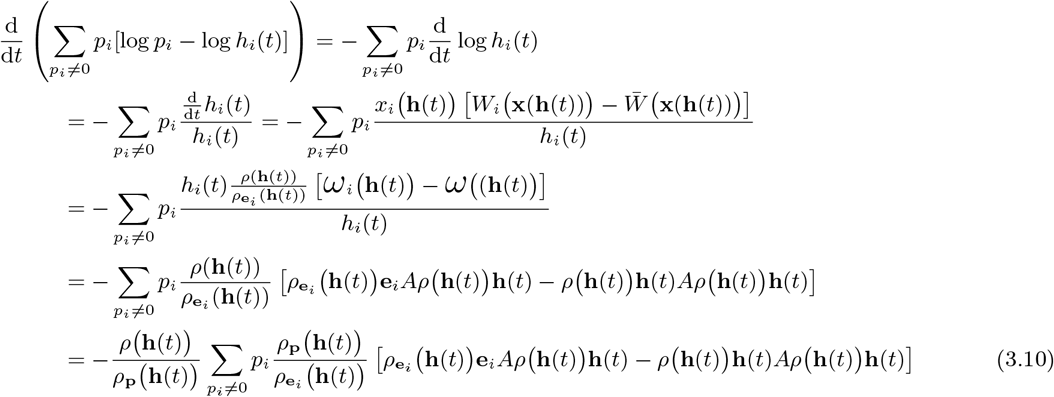

Note that

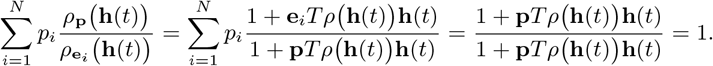

Therefore, (3.10) can be contiued as

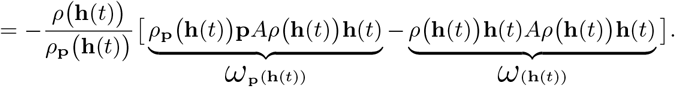

Since **p** is a UESS, by Lemma 2.2, the above expression is strictly negative providing that 0 *<*||**h**(*t*) −**p** ||*< δ* (where *δ* comes from Lemma 2.2). [To avoid interpretation issues (due to the argument of logarithm being 0), it can be assumed that *δ* is already so small that *h*_*i*_(0) *>* 0 (and thus *h*_*i*_(*t*) *>* 0) if *p*_*i*_ *>* 0.]

We conclude, by Lyapunov’s theorem on stability [see for example (Hirsh et al. 2004, p. 194) or (Hofbauer and Sigmund 1998, Theorem 2.6.1)], that **p** is an asymptotically stable rest point of dynamics (3.7).

## 4 Examples

Here, we explore the examples in Varga and Garay (2024) and an example in Varga and Garay (2021) concerning dynamics (3.7). We present the phase portrait for dynamics (3.7) (the larger phase portrait on the left in Figures 1, 2 and 3). Additionally, this phase portrait is transformed by the transformation **h** ↦ **x**(**h**) (appearing as the upper phase portrait on the right in Figures 1, 2 and 3), where 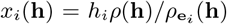 (see Lemma 2.3). The phase portrait with respect to the replicator dynamics (2.2) is also provided to make the comparison easier (the lower phase portrait on the right in Figures 1, 2 and 3). The orbits of dynamics (3.7) depict the evolution of the composition of the subpopulation of active individuals. After the transformation **h** *↦* **x**(**h**), the orbits illustrate the evolution of the composition of the entire population. This should be compared with the orbits for the classical replicator dynamics (2.2).

**Figure 1:**
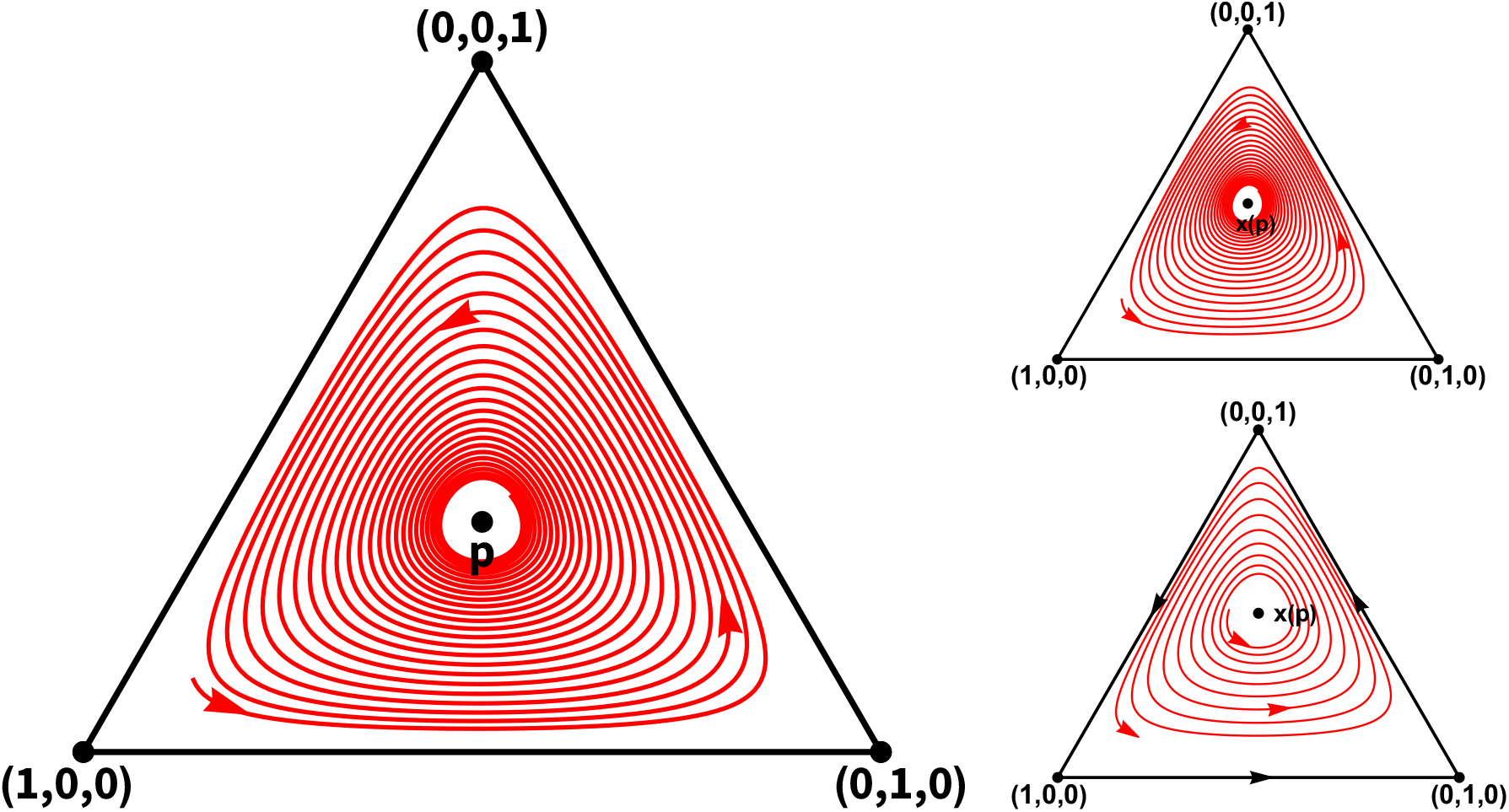
The phase portraits with respect to the matrix game under time constraints with payoff matrix *A* and *T* in (4.11). The *larger diagram on the left* is the phase portrait with respect to dynamics (3.7). Since **p** = (1*/*3, 1*/*3, 1*/*3) is a UESS, it follows that **p** is an asymptotically stable rest point of dynamics (3.7) (see Theorem 3.3). Moreover, the figure suggest the global stability though this has not been proved analytically. In this example, the edges of simplex *S*3 forms a heteroclinic cycle. The *upper diagram on the right* is the transformation of the left-hand side phase portrait by the map **h** *↦* **x**(**h**) (see the paragraph above Definition 2.4). Accordingly, **x**(**p**) is asymptotically stable and the boundary of *S*3 forms a heteroclinic cycle. This diagram should be compared with the *lower phase portrait on the right* which is the phase portrait with respect to the replicator dynamics (2.2). Note that **x**(**p**) is unstable here (for further analyses see Section 3.1 in Varga and Garay 2024).

**Figure 2:**
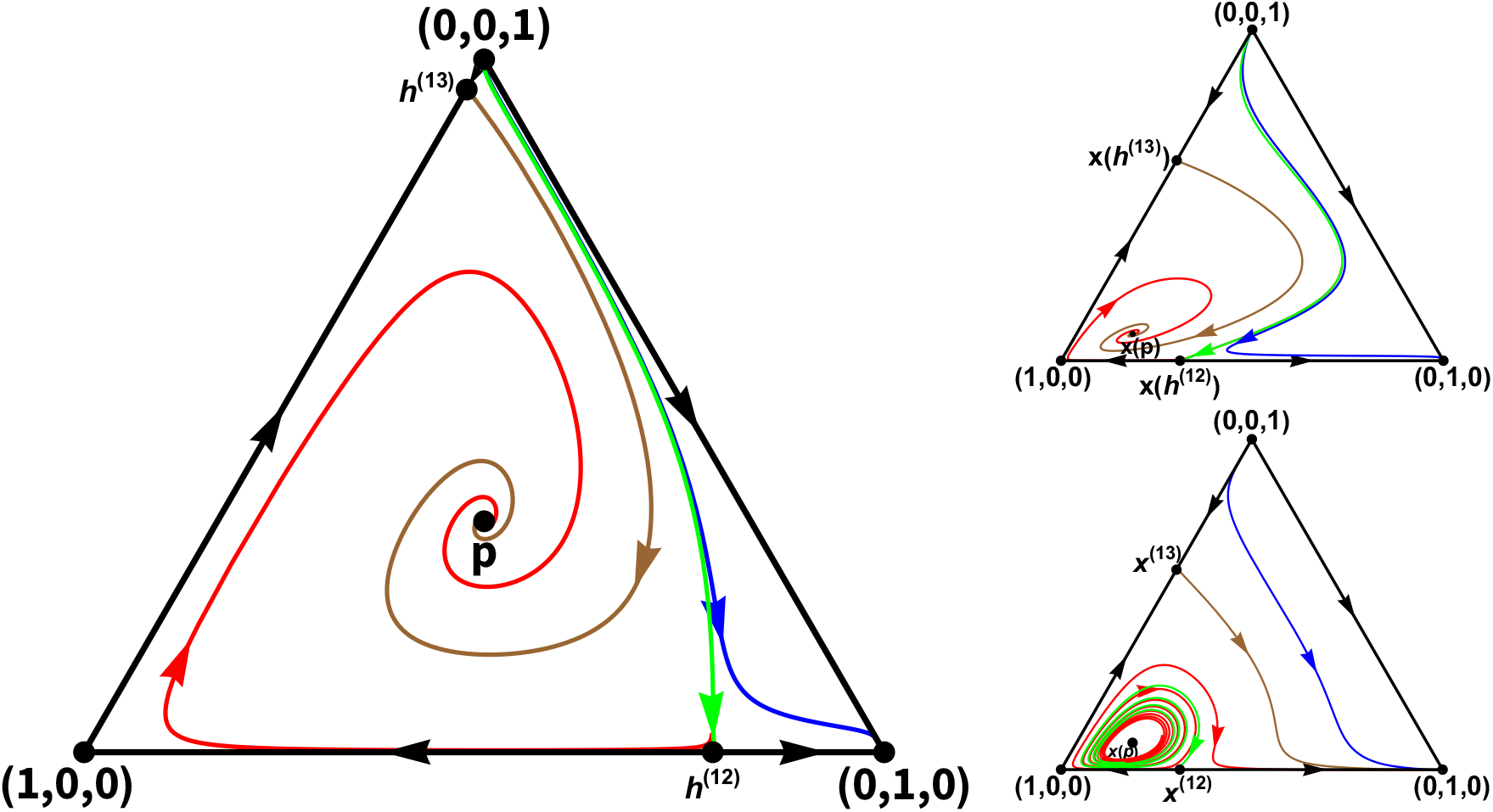
The phase portraits with respect to the matrix game under time constraints with payoff matrix *A* and *T* in (4.12). The *larger diagram on the left* is the phase portrait with respect to dynamics (3.7). Since **p** = (1*/*3, 1*/*3, 1*/*3) is a UESS, it follows that **p** is an asymptotically stable rest point of dynamics (3.7) (see Theorem 3.3). In addition to the vertices, there are two further rest points on the boundary of *S*3. They are **h**^(12)^ on the edge between (1, 0, 0) and (0, 1, 0) and **h**^(13)^ on the edge between (1, 0, 0) and (0, 0, 1), respectively. It seems there is a separatrix starting from (0, 0, 1) and ending in **h**^(12)^ (the green orbit). The orbits on the “left-hand” side of the separatrix in the interior of *S*3 start from (0, 0, 1) except one starting from **h**^(13)^ (the brown orbit) and they all end in **p**. The orbits on the “right-hand” side of the separatrix in the interior of *S*3 also start from (0, 0, 1) but end in (0, 1, 0) which is also asymptotically stable because of (0, 0, 1) being a strict NE (and so a UESS) for matrix game under time constraints (Theorem 4.1 in Varga et al. 2020 and Theorem 3.3). **h**^(12)^ is an unstable rest point on the edge between (1, 0, 0) and (0, 1, 0) while **h**^(13)^ is asymptotically stable on the edge between (1, 0, 0) and (0, 0, 1). The *upper diagram on the right* is the transformation of the left-hand side phase portrait by the map **h** *↦* **x**(**h**) (see the paragraph above Definition 2.4). Accordingly, **x**(**p**) is asymptotically stable. This diagram should be compared with the *lower phase portrait on the right* which is the phase portrait with respect to the replicator dynamics (2.2). Note that **x**(**p**) is unstable here and that a probable separatrix runs from **x**^(13)^ = **x**(**h**^(13)^) to (0, 1, 0). State (0, 0, 1) remains asymptotically stable (Theorem 4.1 in Garay et al. 2018). (For further analyses see Section 3.3 in Varga and Garay 2024.)

**Figure 3:**
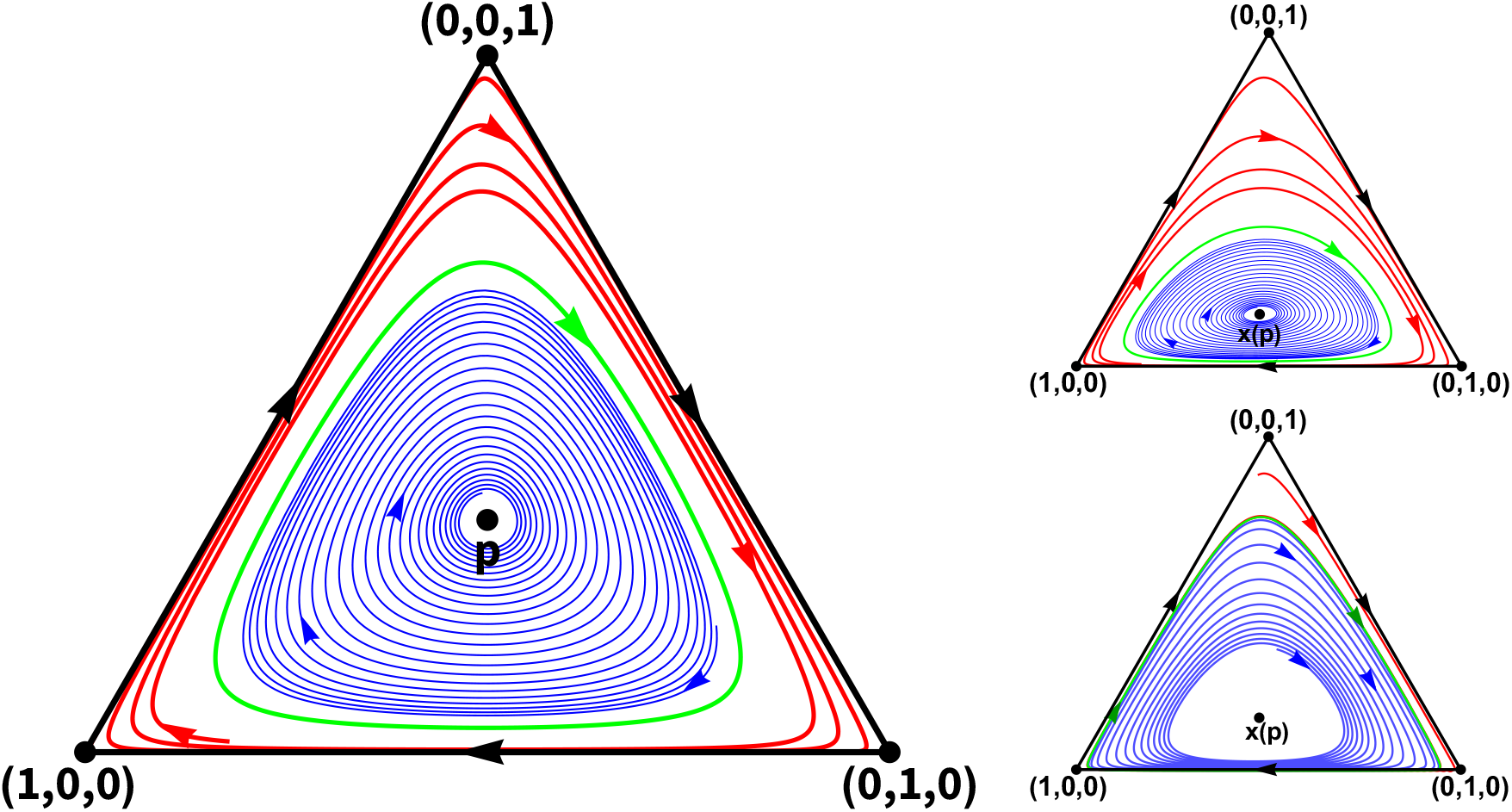
The phase portraits with respect to the matrix game under time constraints with payoff matrix *A* and *T* in (4.13). The *larger diagram on the left* is the phase portrait with respect to the dynamics (3.7). Since **p** = (1*/*3, 1*/*3, 1*/*3) is a UESS, it follows that **p** is an asymptotically stable rest point of dynamics (3.7) (see Theorem 3.3). It seems there is an unstable periodic solution (the green orbit). The orbits outside of the periodic orbit tends to **p** while those outside of the periodic orbit tends to the boundary of *S*3 which forms a heteroclinic cycle. The *upper diagram on the right* is the transformation of the left-hand side phase portrait by the map **h** *↦* **x**(**h**) (see the paragraph above Definition 2.4). Accordingly, **x**(**p**) is asymptotically stable and the boundary of *S*3 forms a heteroclinic cycle. This diagram should be compared with the *lower phase portrait on the right* which is the phase portrait with respect to the replicator dynamics (2.2). Note that **x**(**p**) is unstable here and there is likely a stable periodic orbit (the green orbit) (for further analyses see Section 3.2 in Varga and Garay 2021).

Note that, each example was constructed such that the strategy **p** = (1*/*3, 1*/*3, 1*/*3) is a UESS, but the corresponding state **x**(**p**) is an unstable equilibrium point of the classical replicator dynamics (2.2) (Varga and Garay 2021, Sections A.1 and A.2 in Varga and Garay 2024).

However, we can see that the state **p** = (1*/*3, 1*/*3, 1*/*3) is asymptotically stable for dynamics (3.7), as stated in Theorem 3.3. Consequently, **x**(**p**) is asymptotically stable on the transformed phase portrait, although it is unstable for the replicator dynamics (2.2).

Note that, the dynamics was analytically investigated only on the edges of the simplex *S*_3_ and at the interior rest point **p** = (1*/*3, 1*/*3, 1*/*3) and **x**(**p**), respectively. The phase portrait in the interior of *S*_3_, except of the local neighborhood of **p** and **x**(**p**), respectively, is a result of a numerical simulation using Wolfram Mathematica 12.

### 4.1 Example 1

The payoff matrix *A* and the time constraint matrix *T* are

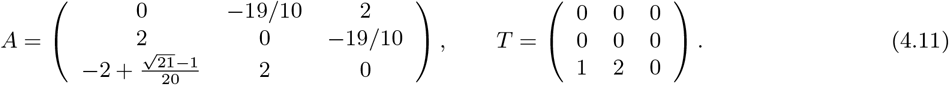

Then **p** = (1*/*3, 1*/*3, 1*/*3) is a UESS (Sections 3.1, A.1 and A.2 in Varga and Garay 2024) and so, by Theorem 3.3, is an asymptotically stable rest point of dynamics (3.7). Under transformation **h** *↦* **x**(**h**), it is transformed into

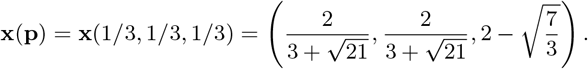

The real parts of the eigenvalues at the rest points for dynamics (3.7) are given in Table 1.

**Table 1:** The real parts of the eigenvalues at the rest points for dynamics (3.7) concerning the matrix game under time constraints with payoff matrix *A* and *T* in (4.11)

| State | Real part of the eigenvalues |  |
| --- | --- | --- |
| (1, 0, 0) | -1.82 | 2 |
| (0, 1, 0) | -1.9 | 2 |
| (0, 0, 1) | -1.9 | 2 |
| $\mathbf{p}$ | -0.01006 | -0.01006 |

The phase portrait with respect to dynamics (3.7) and its transformation by **h** ↦ **x**(**h**), furthermore, the phase portrait with respect to the replicator dynamics (2.2) are shown in Figure 1.

### 4.2 Example 2 with an interior UESS and a strict Nash equilibrium

The payoff matrix *A* and the time constraint matrix *T* are the following in this case:

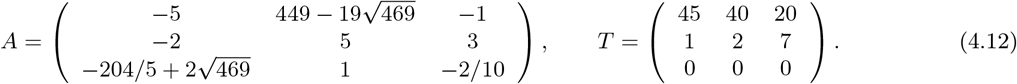

Then **p** = (1*/*3, 1*/*3, 1*/*3) is a UESS (Sections 3.3, A.1 and A.2 in Varga and Garay 2024) and so, by Theorem 3.3, is an asymptotically stable rest point of dynamics (3.7). Under transformation **h** ↦ **x**(**h**), it is transformed into

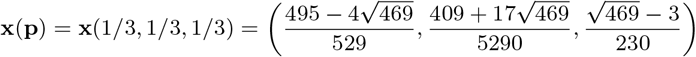

In addition to **p**, pure strategy **e**_2_ = (0, 1, 0) is a UESS too, moreover, a strict NE (Section 3.3 in Varga and Garay 2024), and, due to Theorem 3.3, state (0, 1, 0) is also an asymptotically stable rest point of dynamics (3.7). This is worth comparing with the fact that in the case of classical matrix games, no other Nash equilibrium can exist alongside an internal ESS (the first paragraph after the proof of Theorem 6.4.1 in Hofbauer and Sigmund 1998).

The real parts of the eigenvalues at the rest points for dynamics (3.7) are given in Table 2.

**Table 2:** The real parts of the eigenvalues at the rest points for dynamics (3.7) concerning the matrix game under time constraints with payoff matrix *A* and *T* in (4.12)

| State | Real part of the eigenvalues |  |
| --- | --- | --- |
| $(1, 0, 0)$ | -0.02321 | 0.06136 |
| $(0, 1, 0)$ | -3.743 | -0.375 |
| $(0, 0, 1)$ | 4.6 | 3.2 |
| $\mathbf{p}$ | -0.02237 | -0.02237 |
| $\mathbf{h}^{(12)}$ | -0.07488 | 0.1608 |
| $\mathbf{h}^{(13)}$ | -0.5133 | 1.286 |

The phase portrait with respect to dynamics (3.7) and its transformation by **h** ↦ **x**(**h**), furthermore, the phase portrait with respect to the replicator dynamics (2.2) are shown in Figure 2.

### 4.3 Example 3 with a possible periodic orbit

The payoff matrix *A* and the time constraint matrix *T* are

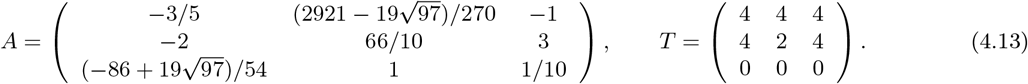

Then **p** = (1*/*3, 1*/*3, 1*/*3) is a UESS (see Section 3.2 in Varga and Garay 2021) and so, by Theorem 3.3, is an asymptotically stable rest point of dynamics (3.7). Under transformation **h** ↦ **x**(**h**), it is transformed into

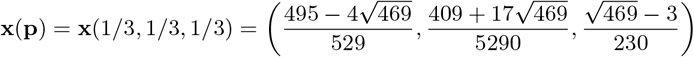

The real parts of the eigenvalues at the rest points for dynamics (3.7) are given in Table 3.

**Table 3:** The real parts of the eigenvalues at the rest points for dynamics (3.7) concerning the matrix game under time constraints with payoff matrix *A* and *T* in (4.13)

| State | Real part of the eigenvalues |  |
| --- | --- | --- |
| $(1, 0, 0)$ | -0.2134 | 0.3211 |
| $(0, 1, 0)$ | -0.575 | 0.05636 |
| $(0, 0, 1)$ | -1.5 | 2.5 |
| $\mathbf{p}$ | -0.004895 | -0.004895 |

The phase portrait with respect to dynamics (3.7) and its transformation **h** ↦ **x**(**h**), furthermore, the phase portrait with respect to replicator dynamics (2.2) are shown in Figure 3. All three phase portraits suggest the existence of an isolated periodic solution (the green orbit), although we have not proven this analytically. Nevertheless, we have demonstrated an example in the case of replicator dynamics where a Hopf bifurcation occurs (Section 3.2 in Varga and Garay 2024), supporting the possibility of an isolated periodic solution under time constraints, even though this is not possible in classical matrix games (Chapter 7.5 in Hofbauer and Sigmund 1998).

## 5 Conclusion

The central relation in classical evolutionary matrix games claims that if **p** is an evolutionarily stable strategy (ESS), then **p** is also an asymptotically stable equilibrium point of the standard replicator dynamics (Hofbauer et al. 1979; Zeeman 1980). This connection is crucial as it links static game theory with a dynamic approach. The game theoretical definition aims to articulate the intuitive biological expectations regarding evolutionary stability into precise mathematical expressions. While the invasion of mutants feels like a dynamic phenomenon, the definition lacks dynamics; it only demands the fulfillment of inequalities to describe that a mutant phenotype cannot invade a population of individuals following an evolutionarily stable strategy. The advantage is that these inequalities are relatively straightforward to test. However, if we want to model not only the evolutionary endpoint of a population, a dynamic model is necessary. One such dynamically relevant model, proposed by Taylor and Jonker (1978), is the replicator dynamics. Although it also allows the examination of the endpoint of the evolution, it requires more complicated mathematical work than simply checking the inequalities.

What is astonishing is that both approaches prove consistent in the sense that a state corresponding to an ESS is an asymptotically stable equilibrium point in the replicator dynamics. This means that the evolutionary endpoint is the same according to both approaches. This consistency is valuable as it allows us to use the simpler static game theory approach instead of the more complex dynamic model.

However, introducing time constraints into the game theoretical model damages this relationship between ESS and replicator dynamics, and generally, it no longer holds. Mathematically, this is not so surprising, as the introduction of time constraints, along with the emergence of new parameters, makes the formulas more complex, providing the possibility for the classical relationship to break down at certain parameter values. Indeed, this is illustrated by examples in Varga and Garay (2021) and Varga and Garay (2024).

This suggests that under time constraints, the replicator dynamics in its classical form may not be the most suitable for mathematically describing intuitive expectations about the evolution of a population. Therefore, in this article, we propose a modification or generalization of the replicator dynamics that aligns better with these intuitive expectations. Considering time constraints, this adjusted dynamics describes the evolution of the ratio of active phenotypes directly, rather than the overall proportion of phenotypes (active and inactive combined), while indirectly determining the latter too. This implies that, under time constraints, the evolution of a population occurs through the active individuals. When there is no time constraint, as in the case of classical evolutionary matrix games, the dynamics coincide with the replicator dynamics. However, when using the generalized dynamics, it still holds true that a state corresponding to a uniformly evolutionarily stable strategy (UESS) is an asymptotically stable equilibrium point of the dynamics.

## Declarations

## Acknowledgements

The author thanks József Garay for his helpful comments on and discussion of this manuscript.

## Funding

TV was supported by project TKP2021-NVA-09. Project no. TKP2021-NVA-09 has been implemented with the support provided by the Ministry of Culture and Innovation of Hungary from the National Research, Development and Innovation Fund, financed under the TKP2021-NVA funding scheme.

This work was supported by University of Szeged Open Access Fund 4680 (to TV).

## Conflict of Interest

The author declares that there are no conflicts of interest regarding the publication of this paper.

## Data availability

The article has no associated data

## Author contributions

Conceptualization: TV. Methodology: TV. Formal analysis: TV. Visualization and numerical investigation: TV. Writing – Review and Editing: TV.

## Footnotes

1 Abusing the notation, we simply write **p***T* **q** and **p***A***q**, respectively, instead of **p***A***q**^*T*^ and **p***A***q**^*T*^, respectively, where the superscript ^*T*^ refers to the transpose of a matrix.

2 Just let *ε* in Definition 2.1 tend to 0 using the continuity of the fitness function in *ε* (Corollary 6.3 in Garay et al. 2018).

3 Note that the notion of ESS equivalent with the notion of UESS in this particular case (Theorem 6.4.1 in Hofbauer and Sigmund 1998).

4 The term “evolutionarily stable state” is used in (Chapter 7.2, Hofbauer and Sigmund 1998) instead of “polymorphic stable state” but we prefer the latter because it may be less confusing.

5 Note that, for classical evolutionary matrix games, *y*_*i*_(**x, p**) = *p*_*i*_ independently of **x** but the independnce of **x** is not sure under time constraints at all.

## Notes

### Competing Interest Statement

The authors have declared no competing interest.

